# Clustered Functional Domains for Curves and Corners in Cortical Area V4

**DOI:** 10.1101/808907

**Authors:** Rundong Jiang, Ian M. Andolina, Ming Li, Shiming Tang

## Abstract

The ventral visual pathway is crucially involved in integrating low-level visual features into complex representations for objects and scenes. At an intermediate stage of the ventral visual pathway, V4 plays a crucial role in supporting this transformation. Many V4 neurons are selective for shape segments like curves and corners, however it remains unclear whether these neurons are organized into clustered functional domains, a structural motif common across other visual cortices. Using two-photon calcium imaging in awake macaques, we confirmed and localized cortical domains selective for curves or corners in V4. Single-cell resolution imaging confirmed that curve or corner selective neurons were spatially clustered into such domains. When tested with hexagonal-segment stimuli, we find that stimulus smoothness is the cardinal difference between curve and corner selectivity in V4. Combining cortical population responses with single neuron analysis, our results reveal that curves and corners are encoded by neurons clustered into functional domains in V4. This functionally-specific population architecture bridges the gap between the early and late cortices of the ventral pathway and may serve to facilitate complex object recognition.

## Introduction

The visual system faces the daunting task of combining highly ambiguous local patterns of contrast into robust, coherent and spatially extensive complex object representations (Connor et al., 2007; Haxby et al., 1991; Mishkin et al., 1983). Such information is predominantly processed along the ventral visual pathway (areas V1, V2, V4, and Inferotemporal cortex [IT]). At early stages of this cortical pathway, neurons are tuned to local single orientation (Hubel and Livingstone, 1987; Hubel and Wiesel, 1968), or combinations of orientations (Anzai et al., 2007; Ito and Komatsu, 2004). Orientation responses are functionally organized into iso-orientation domains that form pinwheel structures in V1 (Ts’O et al., 1990). At later stages like IT, neurons are selective for complex objects predominantly organized categorically (Desimone et al., 1984; Freiwald and Tsao, 2010; Fujita et al., 1992; Kobatake and Tanaka, 1994; Tsao et al., 2003; Tsao et al., 2006). Such complex object organization is embodied using combinations of structurally separated feature columns (Fujita et al., 1992; Rajalingham and DiCarlo, 2019; Tanaka, 2003; Tsunoda et al., 2001; Wang et al., 1996). Positioned in-between the local orientation architecture of V1 and the global object architecture of IT lies cortical area V4, exhibiting visual selectivity that demonstrates integration of simple-towards-complex information (Pasupathy et al., 2019; Roe et al., 2012; Yue et al., 2014), and extensive anatomical connectivity across the visual hierarchy (Gatass et al., 1990; Ungerleider et al., 2008).

Functional organization within V4 has previously been visualized by intrinsic signal optical imaging, and cortical representations of low-level features for orientation, color and spatial frequency have been systematically demonstrated (Conway et al., 2007; Li et al., 2014; Li et al., 2013; Lu et al., 2018; Tanigawa et al., 2010). Such functional clustering suggests that the intracortical organizational motifs in V4 bear some similarity to V1. It remains unknown how more complex feature selective neurons in V4 are spatially organized, and whether feature-like columns found in IT also exist in V4. Because intrinsic imaging is both spatially and temporally limited, it is unable to measure selective responses of single neurons. Using electrophysiology, early studies in V4 using bar and grating stimuli found that V4 neurons are tuned for orientation, size and spatial frequency (Desimone and Schein, 1987). Subsequent studies revealed V4 selectivity for complex gratings and shapes in natural scenes (David et al., 2006; Gallant et al., 1993; Kobatake and Tanaka, 1994). Systematic tuning of V4 neurons for shape segments such as curves and corners as well as combination of these segments was discovered using parametric stimulus sets consisting of complex shape features (Cadieu et al., 2007; Carlson et al., 2011; Oleskiw et al., 2014; Pasupathy and Connor, 1999; Pasupathy and Connor, 2001; Pasupathy and Connor, 2002). Temporally varying heterogeneous fine-scale tuning within the spatial-temporal receptive field has also been observed (Nandy et al., 2016; Nandy et al., 2013; Yau et al., 2013). More recently, artificial neural networks were used to generate complex stimuli that characterize the selectivity of V4 neurons (Bashivan et al., 2019). However, whether such complex-feature selective neurons are spatially organized in V4 remains largely unknown.

In this study, we aim to address whether functional domains encoding complex features such as curves and corners exist in V4. To address this, we performed two-photon (2P) calcium imaging in awake macaque V4. This technique substantially increases the spatial resolution for functional characterization at the single-cell level, while enabling the visualization of spatial distribution and clustering within the cortical population (Garg et al., 2019; Li et al., 2017; Nauhaus et al., 2012; Ohki et al., 2005; Seidemann et al., 2016; Tang et al., 2018). We scanned a large cortical area in dorsal V4 using low-power objective lens to search for regions selectively activated by curves or corners. We subsequently imaged these regions by high-power objective lens to record single neurons’ responses in order to examine whether spatially clustered curve or corner selective neurons could be found. If such neural clusters were found, we further aimed to understand how different curves and corners are encoded and differentiated in greater detail.

## Results

We injected AAV1-hSyn-GCaMP into dorsal V4 (V4d) of two rhesus macaques — GCaMP6f for monkey A and GCaMP5G for monkey B. An imaging window and head posts were implanted 1-2 months after viral injection (see Materials and methods). Subjects were trained to initiate and maintain fixation within a 1° circular window for 2 sec: the first second contained the fixation spot alone, and then stimuli appeared for one second on a LCD monitor positioned 45 cm away (17 inch, 1280 x 960 pixel, 30 pixel/°). Neuronal responses were recorded using 2P calcium imaging, with differential images generated using ΔF = F – F0; where F0 is the average fluorescence 0.5 to 0 sec before stimulus onset, and F is the average response 0.5 to 1.25 sec after stimulus onset.

### Cortical mapping of curve-biased and corner-biased regions in V4

We first identified the retinal eccentricity using drifting gratings for our sites and found they were positioned with an eccentricity of ∼0.7° from the fovea in monkey A and ∼0.9° in monkey B. We next used a low power (4x) objective lens to identify and localized any cortical subregions selectively activated by curves or corners. Using a large range of contour feature stimuli including bars, curves and corners (*Figure 1A*), we scanned a large area (3.4 x 3.4 mm) in V4d (*Figure 1B*), between the lunate sulcus (LS) and the terminal portion of the inferior occipital sulcus (IOS). We obtained global activation maps by Gaussian smoothing (standard deviation σ = 10 pixels, 67 μm) the ΔF/F0 maps. We observed that orientation is organized in linear iso-orientation domains or pinwheel-like patterns, as previously reported (Roe et al., 2012) using intrinsic signal optical imaging in V4 (***Figure S1***).

**Figure 1.**
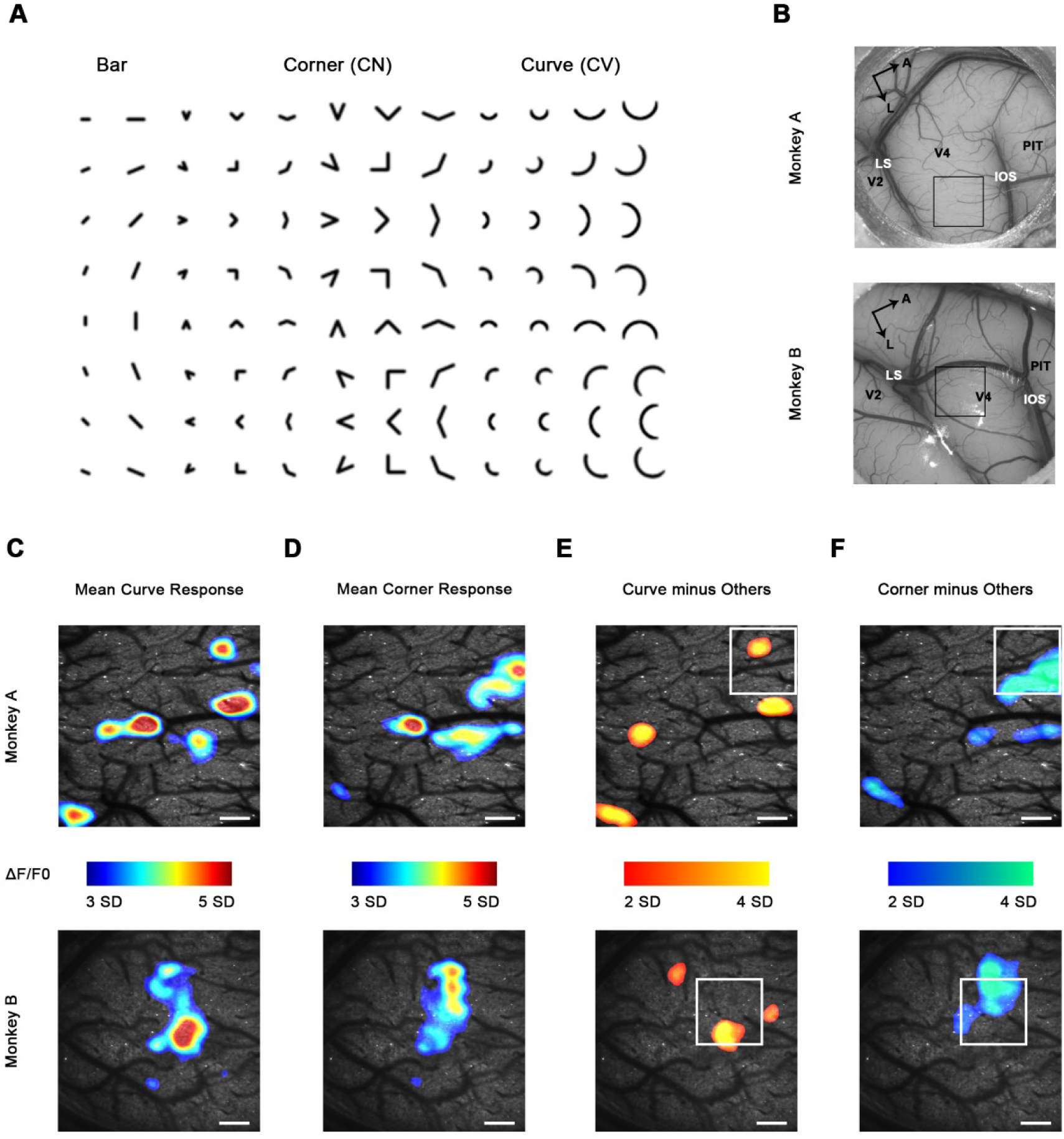
Cortical mapping of curve-biased and corner-biased regions in V4 using a 4x objective lens. (**A**) The stimulus set used for initial cortical mapping consisting of bars, corners and smooth curves. (**B**) Vascular map. LS, lunate sulcus. IOS, inferior occipital sulcus. The black box indicates the imaging site in each subject. (**C**) Cortical regions that showed strong responses (ΔF/F0) to curve stimuli. Scale bar = 400 μm. Threshold = 3 SD. (**D**) Cortical regions that showed strong responses to corner stimuli; same conventions as (C). (**E**) Significant curve-selective regions derived by the average response to curves minus the average response to other stimuli (regions < 400 pixels are excluded). The white box indicates the imaging site selected for 16x objective single-cell mapping. (**F**) Corner selective regions; same conventions as (E).

We then examined the response to curve and corner stimuli in more detail. We averaged the activation maps of all curve stimuli and all corner stimuli respectively, and we found cortical regions responding strongly to curves or corners (*Figure 1C-D*). We computed the response map to curves minus bars and corners to obtain the curve selective activation (*Figure 1E*), revealing several curve selective cortical regions in dorsal V4. We performed significance tests to examine the curve preference (comparing two groups, all curve stimuli against all other stimuli), and all the thresholded regions in Figure 1E passed a one-way ANOVA test at p < 10^−6^ (For monkey A, from left to right, p = 1 × 10^−75^, 2 × 10^−81^, 7 × 10^−66^, 1 × 10^−73^, n = 18; for monkey B, p = 9 × 10^−10^, 4 × 10^−16^, 3 × 10^−12^, n =10). Similarly, we also found regions that were significantly corner selective in dorsal V4 (*Figure 1F*. For monkey A, p = 1 × 10^−44^, 2 × 10^−26^, 1 × 10^−62^, 5 × 10^−26^; for monkey B, p = 2 × 10^−19^). These curve or corner selective regions are considered candidates for functional domains encoding shape segments in V4.

### Single-cell mapping of curve and corner selective neurons reveals they are spatially clustered

To confirm that neurons within these regions were indeed curve or corner selective, we performed single-cell resolution imaging with a high-power objective lens (16×) to record neuronal responses (ΔF/F0) as well as their spatial organization. The imaging sites (850 × 850 μm) in both subjects were chosen to include both curve and corner selective domains found by 4× imaging (*Figure 1E-F*). 535 visually responsive neurons (292 from monkey A and 243 from monkey B) were recorded in total. Each stimulus was repeated 10 times and averaged to derive neuronal responses (***Figure S2***). To characterize neurons’ curve and corner selectivity, we calculated a curve selectivity index (CVSI) and corner selectivity index (CNSI). A Positive CVSI value indicates a neuron’s maximal response to curves is stronger than its maximal response to other stimuli: a CVSI = 0.33 signifies a response twice as strong, and a CVSI = 0.2 is 1.5 times as strong. The same definition applies to CNSI. We found neurons with a high CVSI or CNSI were spatially clustered (*Figure 2A-D*), and these neurons were also selective to the orientation of the integral curves or corners (*Figure 2E-H*; 91.4% of the neurons are significantly tuned to the orientation of curves or corners; one-way ANOVA, Bonferroni corrected p < 0.05). Their overall spatial distribution was consistent with the spatial distribution of curve and corner domains revealed by 4x imaging (*Figure 2A-D vs*. *Figure 1E-F*), especially considering the possible loss of detailed spatial information during Gaussian smoothing of 4x images. This parsimoniously suggests that the observed cortical activation was evoked by responsive neuronal clusters.

**Figure 2.**
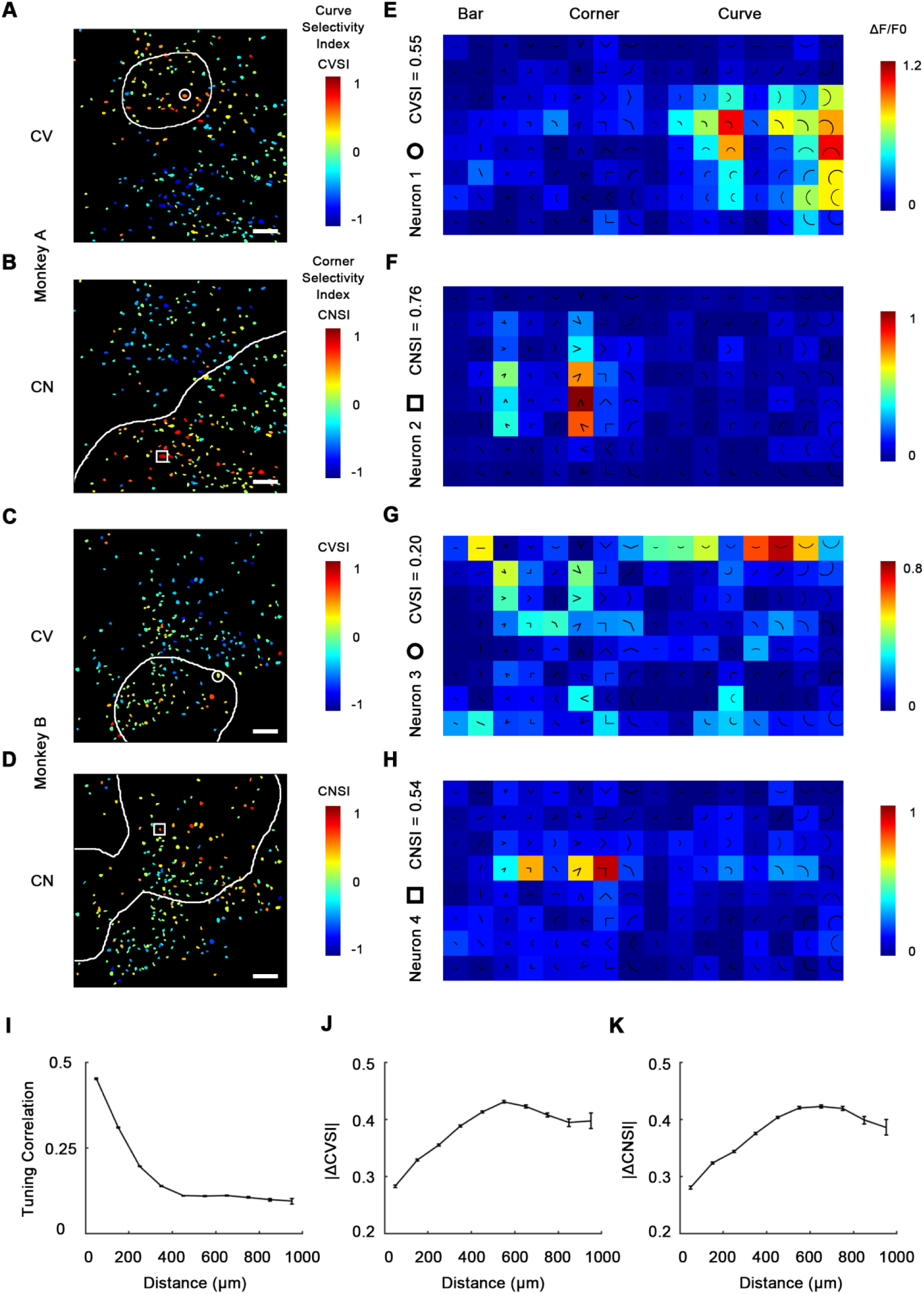
Single-cell mapping of curve and corner selective neurons using a 16x objective lens. (**A**) Cell map of curve (CV) selectivity index (CVSI). Responsive neurons are labeled at their spatial location and colored according to their CVSI. Neurons with high positive CVSI (high curve preference) were clustered in the upper part of the imaging area. The white line indicates the curve-biased regions derived by 4× imaging (Figure 1E). Scale bar = 100 μm. (**B**) Cell map of corner (CN) selectivity index (CNSI). Neurons with high positive CNSI (high corner preference) were clustered in the lower part of the imaging area. (**C**-**D**) Equivalent maps for Monkey B. (**E**-**H**) Responses of 4 example neurons preferring curves or corners, their locations labeled in a-d respectively. (**I**) Neuronal pairwise tuning correlation (mean ± SE) plotted against spatial distances (averaging all neurons every 100 μm). Neurons that are close to each other (< 400 μm) tended to have more similar tuning curves. (**J**) Absolute CVSI value differences (mean ± SE) plotted against distances. (**K**) Absolute CNSI value differences (mean ± SE) plotted against distances.

We next assessed this clustering quantitatively by examining how neuronal responses correlate with spatial distance. For each neuronal pair recorded from the same subject, we computed the pairwise tuning correlation and absolute value differences for CVSI and CNSI plotted against the neuronal pairwise distances. We found that neurons close to each other (< 400 μm approximately) often had more correlated tuning (*Figure 2I*), and generally exhibited more similar CVSI and CNSI values (*Figure 2K-L*). These results indicate curve selective and corner selective neurons are spatially clustered, which could potentially form curve domains and corner domains in V4, which could therefore be detected when imaged at a larger scale.

Out of all 535 neurons recorded from two animals, the majority (346 neurons, 64.7%) significantly preferred curve and corner stimuli over single bars, and only 1.5% (8 neurons) significantly preferred bars over curves and corners (***Figure S3A***), indicating neurons in these areas were indeed much more likely to encode more complex shape features compared to simple orientation. Therefore, we made a combined cell map to depict curve and corner selectivity (*Figure 3A*), neglecting bar responses, by calculating curve/corner index (CVCNI). Similar to CVSI and CNSI, positive CVCNI values indicates a neuron’s maximum response to curves is stronger than its maximum response to corners, and vice versa. As expected, neurons with similar CVCNI values were spatially clustered. Neurons that fell into the 4x-defined curve domains generally had positive CVCNI values (*Figure 3B*), and those in the 4x corner domains generally had negative CVCNI values (*Figure 3C*). We also performed a one-way ANOVA comparing neurons’ maximum curve and corner responses. We found neurons with CVCNI > 0.2 or < −0.2 (which means 1.5 times as strong) predominantly showed significant preferences (p < 0.05) to curves or corners over the other kind (*Figure 3D*). The curve or corner selective neurons (red and blue neurons in *Figure 3D*) have very diverse curve or corner tuning, and could be either selective or invariant to the radius and radian of curves or bar length and separation angle of corners (***Figure S3B-D***), which potentially enables the encoding of multiple shape segments. More interestingly, these neurons that were heavily biased to curves or corners over the other tended to respond very weakly to single bars (*Figure 3E*), implying they might be detecting more complex and integral shape features instead of local orientation. These results suggest that curves and corners are encoded by different neuronal clusters organized in curve and corner domains, and these domains are distinct from those representing single orientations.

**Figure 3.**
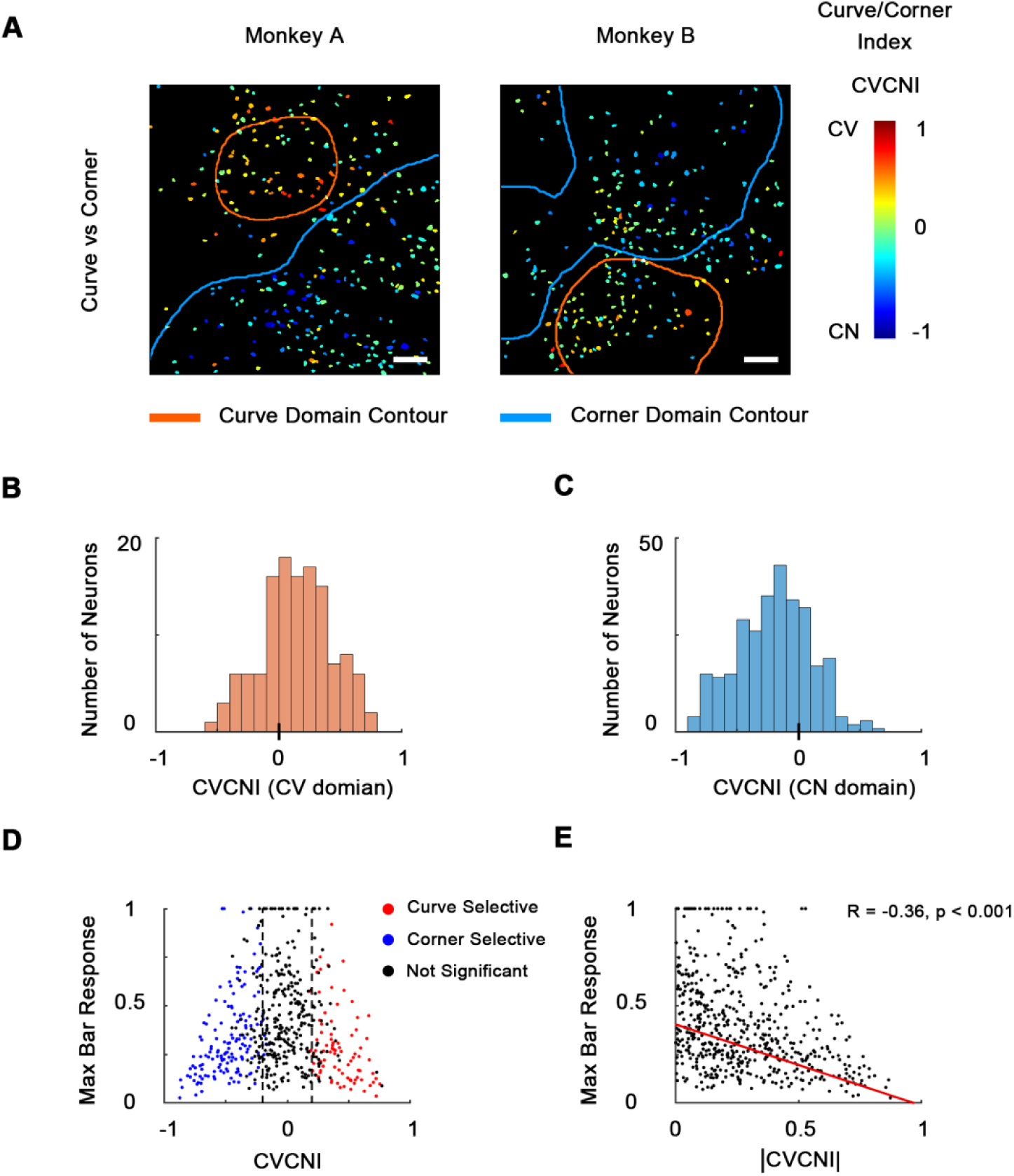
Combined maps of curve/corner preference. (**A**) Cell map of curve/corner index (CVCNI). Positive CVCNI indicates preference for curves over corners and vice versa. Curve selective neurons and corner selective neurons are spatially clustered. Scale bar = 100 μm. (**B**) Histogram of CVCNI for neurons located within the curve-biased domains. Mean = 0.15 ± 0.03 S.E. (**C**) Histogram of CVCNI for neurons located within the corner-biased domains. Mean = −0.20 ± 0.02 S.E. (**D**) Scatterplot of maximum responses to bars (normalized to 0-1 by the maximum responses to all contour features) against CVCNI. Red dots indicate neurons showing significant preference for curves (ANOVA p < 0.05, n = 10) and blue for corners. The majority of neurons (74.5%) with CVCNI < −0.2 or > 0.2 were significantly selective. Neurons that highly preferred curves over corners or corners over curves didn’t respond strongly to single orientated bars. (**E**) Neurons’ maximum bar responses were negatively-correlated with the absolute values of CVCNI. The red line represents the linear regression line.

### Curve preferring neurons are selective for smoothness

Curves and corners are both different from single bars in that they potentially contain multiple different local orientations, yet we found them to be encoded by different neuronal clusters in V4. This suggests that V4 neurons are not recognizing shapes with more than one local orientation, but computing a more fundamental feature difference. To investigate what distinguishes curves from corners in V4, we tested hexagonal segments (Π-shape stimuli; *Figure 4A*) that highly resemble curves except for a lack of smoothness (Nandy et al., 2013). We found that neurons that were very selective to smooth curves did not respond strongly to Π-shape stimuli (*Figure 4A*), suggesting they were selective to smoothness, rather than multiple orientations. In the same way as CVCNI, we calculated CVPII, which characterizes a neuron’s preference to smooth curves over the Π-shape stimuli. We found neurons’ CVPII were highly consistent with CVCNI (R = 0.72, p < 0.001, *Figure 4B*), which means neurons preferring smooth curves over corners would also prefer smooth curves over Π-shape stimuli. As a result, the maps of CVPII were also consistent to CVCNI maps (*Figure 4C vs*. *Figure 3A*). K-means clustering analysis of population responses also showed that smooth curves are encoded differently from rectilinear shapes including Π-shapes and corners (***Figure S4***). Therefore, smoothness is important to the distinct encoding of curves and corners in the specific curve domains and corner domains in V4.

**Figure 4.**
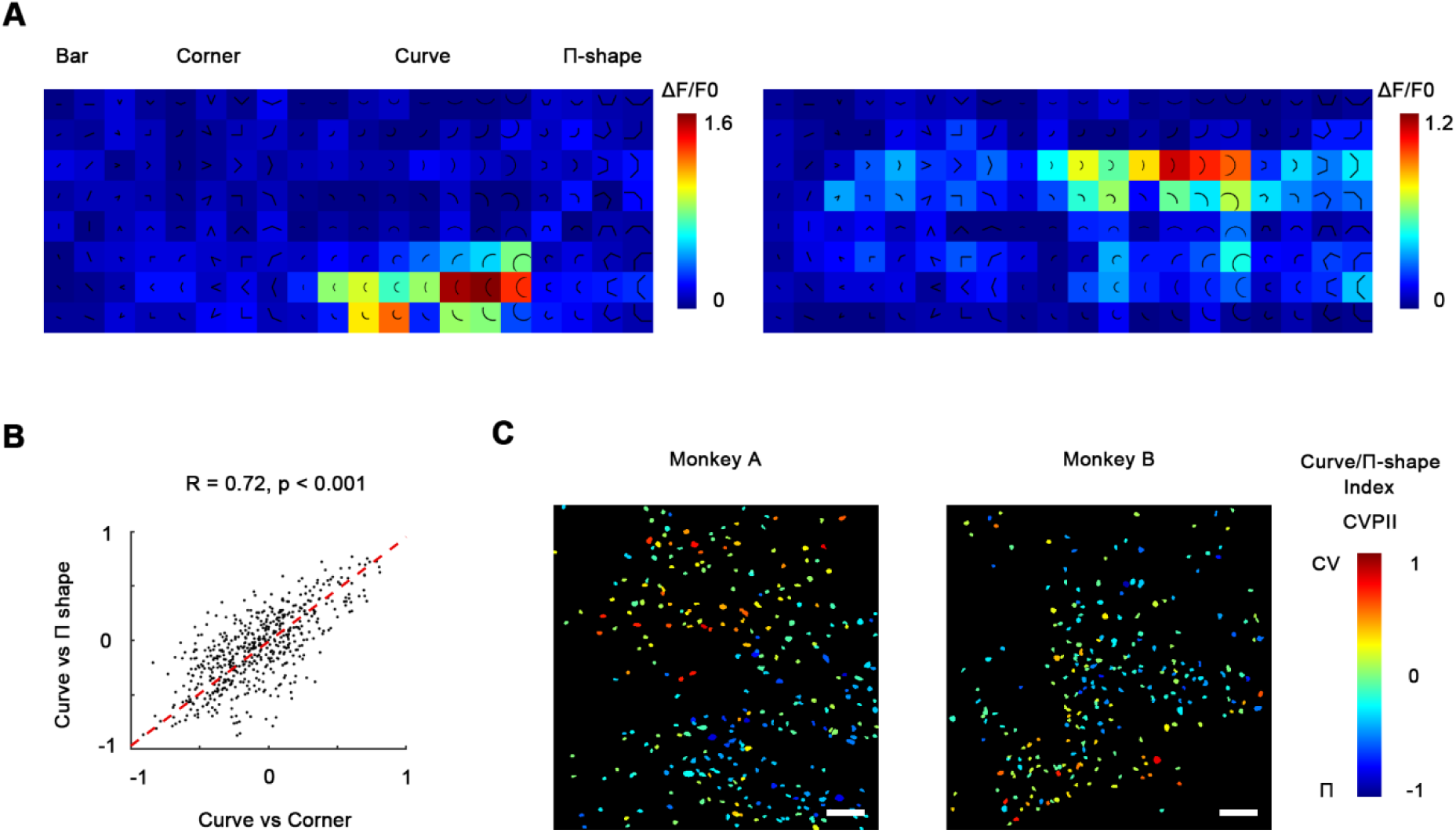
Curve preferring neurons are selective for smoothness. (**A**) Responses of 2 example curve preferring neurons to bars, corners, smooth curves and Π-shape stimuli. The neurons responded strongly to smooth curves but not to Π-shape, which highly resemble curves despite lack of smoothness. (**B**) Scatterplot of CVCNI against CVPII, which characterizes neuronal preference for smooth curves over Π-shape stimuli. The red dash line represents the linear regression line. The two values were highly correlated, indicating neurons preferring curves over corners also preferred curves over Π-shape stimuli. Scale bar = 100 μm. (**C**) Cell map of CVPII. Neurons are clustered similarly to CVCNI (Figure 3A).

### Curve and corner selectivity is related to concentric and radial grating preference

Early studies in V4 demonstrated that many V4 neurons are selective for non-Cartesian gratings (David et al., 2006; Gallant et al., 1993). Concentric gratings highly resemble curves, while radial gratings resemble corners, and we wondered whether these two types of gratings are also separately encoded by neurons in curve and corner domains. So in addition to contour feature stimuli, we also tested concentric, radial and Cartesian gratings (***Figure S5A***). The resultant selectivity maps were consistent with the contour feature maps as predicted. 48.4% of the neurons recorded in the imaging areas significantly preferred concentric or radial gratings over Cartesian gratings, while only 2.2% significantly preferred Cartesian gratings (***Figure S5B***). In addition, many of them were heavily biased to one over the other. Similar to CVCNI, we computed concentric/radial index (CRI) to characterize this bias. CRI and CVCNI values were found to be correlated (R = 0.38, p < 0.001, *Figure 5B*), and naturally their cell maps were also consistent (*Figure 5C vs*. *Figure 3A*), suggesting that classical polar grating selectivity is closely related to curve and corner selectivity. Meanwhile, to assess whether the observed selectivity is related to different spatial frequencies, we examined the CRI map at 1, 2 & 4 cycle/°. The CRI values of all neurons at 3 spatial frequencies are highly correlated (Pearson correlation, all R > 0.5, p < 0.001), and the map structures were found to remain consistent across 3 spatial frequencies (*Figure 5C*), implying such selectivity is not directly related to spatial frequency.

**Figure 5.**
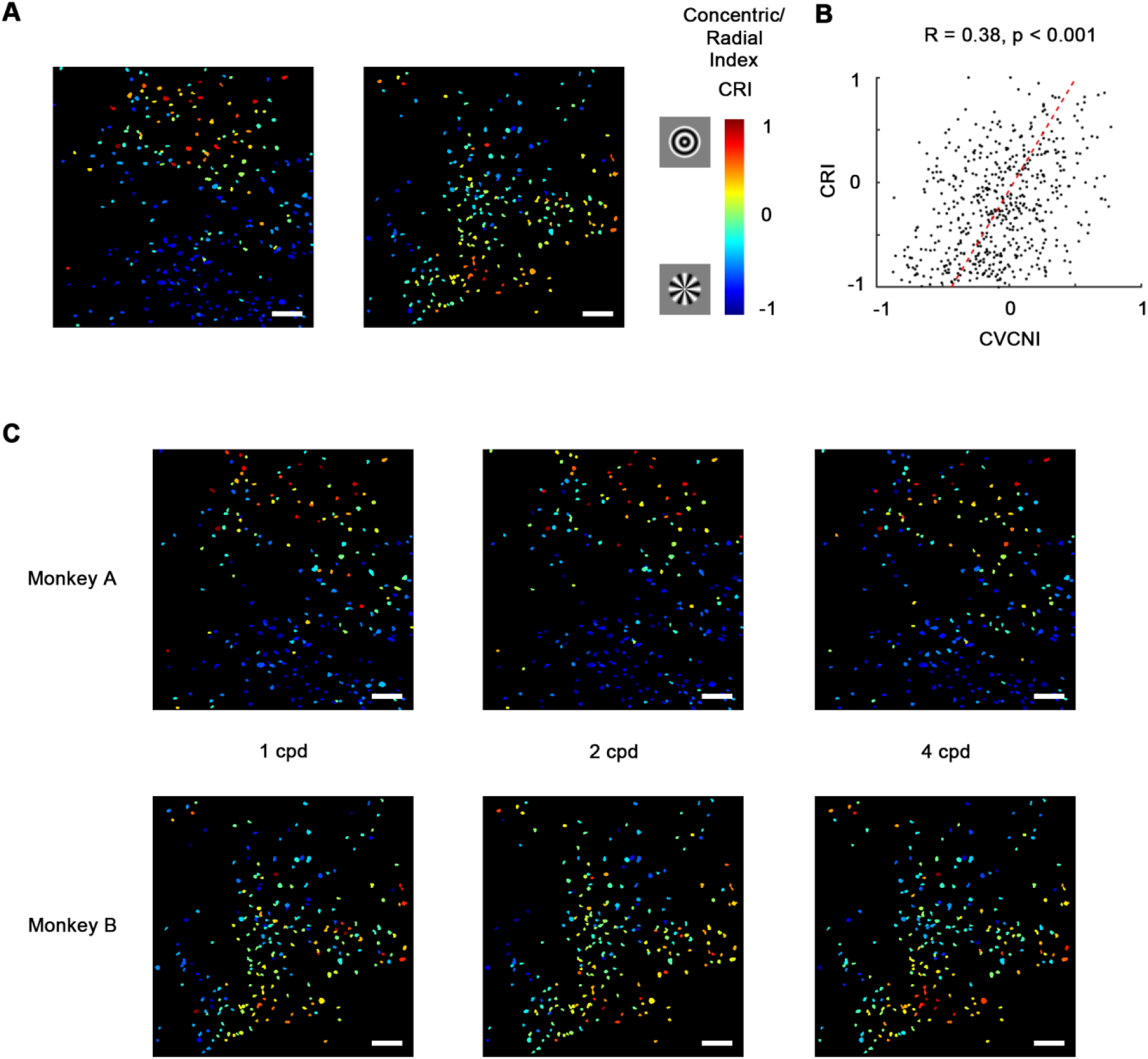
Concentric and radial gratings preference. (**A**) Cell map of concentric/radial index (CRI). Positive CRI indicates preference for concentric over radial gratings and vice versa. Concentric grating selective neurons and Radial grating selective neurons are spatially clustered, and the overall distribution was consistent with curve/corner selectivity (Figure 3A). Scale bar = 100 μm. (**B**) Scatterplot of CVCNI against CRI, which were positively correlated. The red dash line represents the linear regression line. (**C**) CRI cell maps at spatial frequencies of 1, 2, 4 cycle/° (cpd). The map structure remained consistent.

## Discussion

Using two-photon Calcium imaging, we identified cortical subregions in macaque V4d selective for curves or corners (*Figure 1E-F*), with individual curve and corner selective neurons consistently clustered spatially (*Figure 3A*). These neurons exhibited diverse curve or corner selectivity (***Figure S3B-D***) and could potentially be involved in the encoding and processing of a large variety of curves and corners. These results demonstrate the existence of functionally-specific curve and corner domains in V4d.

Functional organization for low-order orientation and spatial frequency representations in macaque V4 had previously been visualized using intrinsic signal optical imaging (Lu et al., 2018; Roe et al., 2012). For more complex-shape features, very few studies have been carried out in V4 to characterize its functional organization, let alone at single-cell resolution. We report here the existence of cortical micro-domains consisting almost entirely of neurons selective for curves. This finding at the single-cell level is consistent with an fMRI study which reported curvature-biased patches in macaque V4 (Yue et al., 2014). The patches we found were smaller in size (about 400 μm) than those observed using fMRI; we suspect due to the improved spatial resolution afforded by two-photon imaging. Additionally, we found cortical domains in V4d selective for corners. To the best of our knowledge, this is the first time the combined functional organization of smooth curves and rectilinear corners in V4 has been demonstrated at single-cell resolution.

A number of electrophysiology studies have reported that some neurons in V4d are selective for more complex features (Gallant et al., 1993; Hegde and Van Essen, 2007; Kobatake and Tanaka, 1994; Pasupathy and Connor, 1999). Our results, consistent with these works, identified many curve or corner selective neurons. In addition, given the ability of 2P-imaging to quantify the spatial relationships between neurons, we further revealed that they are spatially clustered.

We observed some deviations of our results from earlier studies. First, the percentage of complex feature selective neurons we found in our study is higher than previously observed (Gallant et al., 1993; Pasupathy and Connor, 1999); in our hands the vast majority neurons preferred curves and corners over bars and concentric and radial gratings over Cartesian gratings. Second, it was previously reported that curvature selective neurons in V4 also respond to rectilinear Π-shapes (Nandy et al., 2013). This was not the case for our data, which we infer is primarily due to sampling neurons within or close to curve and corner domains (which is difficult to detect with classical electrophysiology). We do not wish to infer that curve and corner stimuli are only encoded by neurons in the curve and corner domains while other neurons are not involved. But we demonstrated neurons in the curve and corner domains are tuned to more complex and integral features rather than local orientation or spatial frequency, alone, supporting the encoding of shape segments with intermediate complexity in V4 (Bushnell and Pasupathy, 2012; El-Shamayleh and Pasupathy, 2016; Oleskiw et al., 2014; Rust and Dicarlo, 2010).

Complexity increases as visual shape information is processed along the ventral visual pathway. Neurons in V1 are tuned to low-order orientation and spatial frequency and organized in iso-orientation domains and orientation pinwheels (Nauhaus et al., 2012; Ts’O et al., 1990). Neurons in IT are selective for complex features and objects and organized in feature columns and face patches (Tanaka, 2003; Tsao et al., 2006; Tsunoda et al., 2001). The simple-to-complex transformation and integration take place in the intermediate stages between V1 and IT. Researchers have reported that some V2 neurons are selective to combination of multiple local orientations, from which corner selectivity might emerge (Anzai et al., 2007; Ito and Komatsu, 2004). Our results in V4d showed that intermediate shape segments like curves and corners are separately encoded by neurons in specific functional domains, and the curve and corner selective neurons are tuned to the integral features instead of local orientation. It is possible that these complex feature-selective neurons receive inputs or modulation from nearby orientation neurons, which might underlie previous findings that the response profiles of V4 neurons were temporally heterogeneous (Nandy et al., 2016). Given that we record neurons whose stimulation is not isolated to the ―optimal‖ spatial location in the receptive fields (i.e. the RF locations of some neurons might deviate for the population RF), the nature of the domains may also be modulated by stimulus translation variance, and future studies addressing positional variance and stimulus encoding are warranted. Our sample of V4d was also near-foveal in terms of eccentricity. It is well established that the ventral pathway connectivity to IT favors central rather than peripheral visual space (Ungerleider et al., 2008), but the relationship of visual eccentricity to these functional domains remains unknown. The existence of curve and corner domains for neuronal encoding in V4d provides significant support for integration of shape information in the intermediate stages of the visual hierarchy. These findings provide more comprehensive understanding of the functional architecture of V4 feature selectivity.

Finally, our results may also help to explore the later stage of the visual hierarchy. The data suggests that higher-order pattern domains may emerge gradually along the ventral pathway. The curve and corner domain responses in V4 could possibly form the basis for more complex feature columns, object domains and even face patches in IT. Studying such functional cross-areal connectivity remains an critical goal for future studies of the visual system. It is also interesting to try to identify why smooth curves and rectilinear corners are separated as early as V4. One possible explanation is that in natural environments, smooth curves are more prevalent in *e.g.* living animals or foods that are of particular interest to primates, while corners are often found in the background environment of stones or branches (i.e. such differences underlie statistical regularities in natural images of objects). The separation of curves and corners in V4 might facilitate the statistical classification of visual object categories in IT or higher cortices. Such comparisons will provide a basis for future investigations that can compare the statistical feature relationships for natural images between V4 and IT functional domains.

## Materials and methods

All procedures involving animals were in accordance with the Guide of Institutional Animal Care and Use Committee (IACUC) of Peking University Laboratory Animal Center, and approved by the Peking University Animal Care and Use Committee (LSC-TangSM-5).

### Animal preparation

The subjects used in this study were two adult male rhesus monkeys (*Macaca mulatta*, 4 and 5 years of age, respectively), purchased from Beijing Prima Biotech Inc. and housed at Peking University Laboratory Animal Center. Two sequential surgeries were performed on each animal under general anesthesia. In the first surgery, we performed a craniotomy over V4 and opened the dura. We injected 200 nl of AAV9.Syn.GCaMP6f.WPRE.SV40 (CS1001, titer 7.748e13 [GC/ml], Penn Vector Core) or AAV1.Syn.GCaMP5G.WPRE.SV40 (V4102MI-R, titer 2.37e13 [GC/ml], Penn Vector Core) at a depth of about 350 μm. Injection and surgical protocols followed our previous study (Li et al., 2017). After injections, we sutured the dura, replaced the skull cap with titanium screws, and closed the scalp. The animal was then returned for recovery and received Ceftriaxone sodium antibiotic (Youcare Pharmaceutical Group Co. Ltd., China) for one week. 45 days later, we performed the second surgery to implant the imaging window and head posts. The dura was removed and a glass coverslip was put directly above the cortex without any artificial dura and glued to a titanium ring. We then glued the titanium ring to the skull using dental acrylic. The detailed design of the chamber and head posts can be found in our previous study (Li et al., 2017).

### Behavioral task

Monkeys were trained to maintain fixation on a small white spot (0.1°) while seated in a primate chair with head restraint to obtain a juice reward. Eye positions were monitored by an ISCAN ETL-200 infrared eye-tracking system (ISCAN Inc., Woburn, MA) at a 120 Hz sampling rate. Trials in which the eye position deviated 1° or more from the fixation point were terminated and the same condition was repeated immediately. Only data from the successful trials was used.

### Visual stimuli

The visual stimuli were displayed on an LCD monitor 45 cm from the animal’s eyes (Acer v173Db, 17 inch, 1280 x 960 pixel, 30 pixel/°, 80 Hz refresh rate). After acquiring fixation, only the grey background was presented for the first 1 sec to obtain the fluorescence baseline, and then the visual stimuli were displayed for further 1 sec. No inter-trial interval was used. Stimuli were presented in pseudo-random order. We used square-wave drifting gratings (0.4° diameter circular patch, full contrast, 4 cycle/°, 3 cycle/sec) generated and presented by the ViSaGe system (Cambridge Research Systems, Rochester, UK) to measure the retinal eccentricity, which was about 1° bottom left to the fovea for both monkey.

Contour feature stimuli were generated using Matlab (The MathWorks, Natick, MA) and presented using the ViSaGe system (Cambridge Research Systems, Rochester, UK). The contour feature stimuli were 2 pixels wide. The lengths of the bars and corner edges were 10 and 20 pixels (30 pixel/°, 0.33° and 0.67°), and the radius of curve stimuli were also 10 and 20 pixels. For each of the 2 sizes, the curve stimuli varied in radians (120°, 180° for 4x imaging and 60°, 90°, 120°, 180° for 16x imaging). The corner stimuli also varied in 3 separation angles (45° and 90° and 135°). All contour feature stimuli were rotated to 8 orientations (0°, 45°, 90°, 135°, 180°, 225°, 270°, 315° for curves and corners, and 0°, 22.5°, 45°, 67.5°, 90°, 112.5°, 135°, 157.5° for bars).

The Cartesian (8 orientations, 0°, 45°, 90°, 135°, 180°, 225°, 270°, 315°), concentric and radial grating stimuli were full contrast sinusoidal gratings (edge blurred) which were 90 pixels (3°) in diameter, with spatial frequencies (SF) of 1, 2, 4 cycle/°. The concentric gratings were generated as

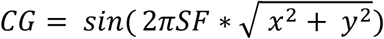

The radial gratings were generated as

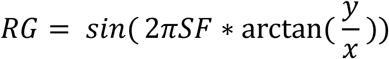

### Two-photon imaging

Two-photon imaging was performed using a Prairie Ultima IV 2P laser scanning microscope (Bruker Corporation, Billerica, MA) during experiments. 1000 nm mode-lock laser (Spectra-Physics, Santa Clara, CA) was used for excitation of GCaMPs, and resonant galvo scanning (512 x 512 pixel, 32 frame/sec) was used to record the fluorescence images (8 fps, averaging every 4 frames). A 4x objective (Nikon Corporation, Tokyo, Japan) was used for sub-cortical level recording (3.4 x 3.4 mm, 6.7 μm/pixel), and a 16x objective (Nikon Corporation, Tokyo, Japan) for neural population recording at single-cellular resolution (850 x 850 μm, 1.7 μm/pixel). We used a Neural Signal Processor (Cerebus system, Blackrock Microsystem, Salt Lake City, UT) to record the time stamp of each frame of the 2P images as well as the time stamps of visual stimuli onset for synchronization.

### Image data processing

Image data was processed by Matlab. The two-photon images were first aligned to a template images by a 2D cross-correlation algorithm (Li et al., 2017) to eliminate motion artifacts during recording sections. For all the successful trials, we found the corresponding 2P images by synchronizing the time stamps of stimulus onset recorded by the Neural Signal Processor (Cerebus system, Blackrock Microsystem, Salt Lake City, UT). The differential fluorescence image was calculated as ΔF = F – F0, where the basal fluorescence image F0 was defined as the average image of 0-0.5 sec before stimulus onset, and F as the average of 0.5-1.25 sec after stimulus onset, both averaged across all repeats for each stimulus.

For 4x imaging, the ΔF/F0 maps were Gaussian smoothed using a low-pass Gaussian filter (σ = 10 pixels) to obtain the activation maps. For 16x imaging, to identify responding cell bodies (ROIs), the differential images (ΔF) went through a band-pass Gaussian filter (σ = 2 pixels and 5 pixels respectively, only used for identifying ROIs) and were then binarized using a pixel value threshold of 3 SD. The connected components (>25 pixels) were identified as candidates for active ROIs. The roundness of these ROIs was calculated as

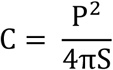

P is the perimeter the ROI and S is the area. Only ROIs with C < 1.1 were identified as cell bodies.

### Curve and corner domains

The curve selective activation map was generated by all curves minus all others, which is the average smoothed ΔF/F0 map of all 32 curves minus that of all 64 other stimuli (48 corners and 16 bars). The regions above 3 SD (>= 400 pixles) were then tested by one-way ANOVA comparing the responses to all the 32 curve stimuli and all the 64 other stimuli (comparing these two groups, repeats n = 18 and 10 for the two animals respectively). All the thresholded regions in Figure 1E-F passed the test with significance p < 10^−6^. Multiple comparisons (comparing the 96 stimuli) were also performed with Bonferroni corrected p < 0.001. The corner selective activation followed the same procedure. The contours of curve and corner domains in 4x imaging were directly copied to 16x after aligning the reference images.

### Quantification and statistical analysis

Curve Selectivity Index (CVSI) is used to characterize a neuron’s preference to curves over other stimuli (bars and corners), defined as

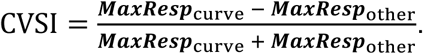

***MaxResp***_curve_ is the neuron’s maximum response to curve stimuli and ***MaxResp***_other_ is the neuron’s maximum response to other stimuli (bars and corners). CVSI ranges from −1 to 1, and positive CVSI value indicates a neuron’s response to its optimal curve stimuli is greater than its response to optimal bar or corner stimuli.

Corner Selectivity Index (CNSI) is defined as

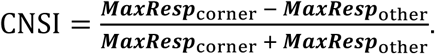

***MaxResp***_corner_ is the neuron’s maximum response to corner stimuli and ***MaxResp***_other_ is the neuron’s maximum response to other stimuli (bars and curves).

Curve/Corner Index (CVCNI) is defined as

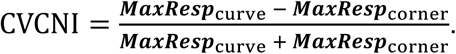

We also performed one-way ANOVA test comparing neuron’s maximum response to curve stimuli and maximum response to corner stimuli in Figure 3D, with threshold value p = 0.05, repeats n = 10.

Curve/Π-shape Index (CVPII) is defined as

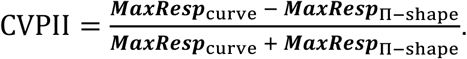

***MaxResp***_Π-shape_ is the neuron’s maximum response to Π-shape stimuli. The Pearson correlation of CVCNI and CVPII was calculated in Figure 4B, and the regression line was derived by minimizing 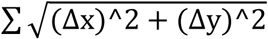.

Concentric/Radial Index (CRI) is defined as

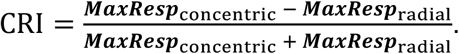

***MaxResp***_concentric_ is the neuron’s maximum response to concentric gratings and ***MaxResp***_radial_ is the neuron’s maximum response to radial gratings. The Pearson correlation of CVCNI and CRI was also calculated in *Figure 5B*, and the regression line was derived by minimizing 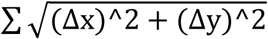.

### Clustering analysis

We analyzed 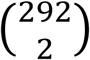 neuron pairs from monkey A and 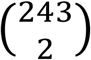 neuron pairs from monkey B in *Figure 2I-K*. Pairwise tuning correlation was calculated as the Pearson correlation of the two neurons’ responses to all bar, curve and corner stimuli, and were plotted against pairwise spatial distances (averaging all neurons every 100 μm). Similarly, the differences in CVSI and CNSI were also plotted against pairwise spatial distances.

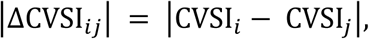

where CVSI_*i*_ is the CVSI of neuron *i* and CVSI_*j*_ is the CVSI of neuron *j*.

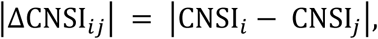

where CNSI_*i*_ is the CVSI of neuron *i* and CNSI_*j*_ is the CNSI of neuron *j*.

### K-means analysis

We performed K-means analysis to cluster the stimulus forms and the neurons. Responses of 535 neurons to 20 forms (2 bars, 8 curves, 6 corners, and 4 Π-shapes, each at 8 orientations) are used to construct the responses matrix R as

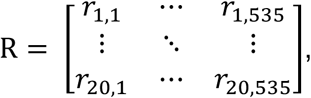

where *r*_*i,j*_ is the response of neuron *j* to stimulus form *i*. Only the maximum responses among 8 orientations were used, as 488 out of all 535 neurons (91.2%) are significantly (one-way ANOVA, Bonferroni corrected p < 0.05) tuned to the orientation of the optimal form.

We used population response vectors (*RP,* rows of matrix R) to cluster the forms. For form *i*,

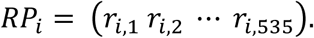

We used neuron response vectors (*RN*, columns of matrix R) to cluster the neurons. For neuron *j*,

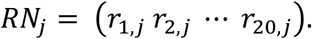

The number of clusters was determined using Calinski-Harabasz criterion and squared Euclidean distance. Maximum literation time = 10000. Clustering was repeated for 10000 times with new initial cluster centroid, and the one with the lowest within-cluster sum was used.

### Multi-dimensional scaling

Classical multi-dimensional scaling (MDS) was performed to visualize the clustering of stimulus forms derived by K-means. The distance (dissimilarity matrix) was computed as

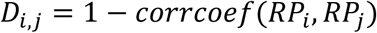

*D_i,j_* is the distance between form *i* and *j*, *corrcoef* is the Pearson correlation, and *RP_i_* is the population response vectors of form *i*. Classical multi-dimensional scaling was performed using singular value decomposition (SVD) algorithm.

The normalized stress was computed as

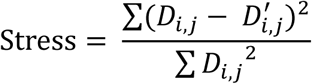

*D*_*i,j*_ is the distance in the original space and *D*’ is the distance in the new MDS space.

### Data and code availability

All data and MATLAB codes are available upon request.

## Acknowledgements

This work was supported by National Natural Science Foundation of China (grant no. 31730109), National Basic Research Program of China (grant no. 2017YFA0105201), National Natural Science Foundation of China Outstanding Young Researcher Award (grant no. 30525016) and a Project 985 grant of Peking University, Beijing Municipal Commission of Science and Technology (grant no. Z151100000915070).

## Author contributions

Author contributions: S.T. and R.J. conceived the project and designed the experiments; R.J. and M.L. performed the experiments; R.J. and I.M.A. analyzed the data and wrote the manuscript; S.T. supervised the project.

