## Supplementary material for "Clustered Functional Domains for Curves and Corners in Cortical Area V4": Figure S

### Supplementary Materials

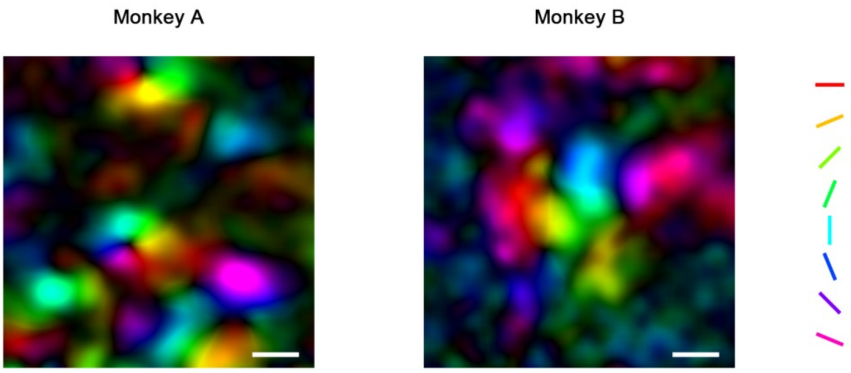


**Figure S1. Related to Figure 1.** Pseudo-color orientation map obtained by 4× imaging. Scale bar = 400 μm. For each pixel, the preferred orientation was derived by the vector summation of its responses (Gaussian smoothed ΔF/F0) to 8 orientations. Hues in the map indicate the orientations of the sum vectors (preferred orientation), and lightness indicates the length (orientation selectivity). Orientation is organized in linear bands and pinwheels in V4.

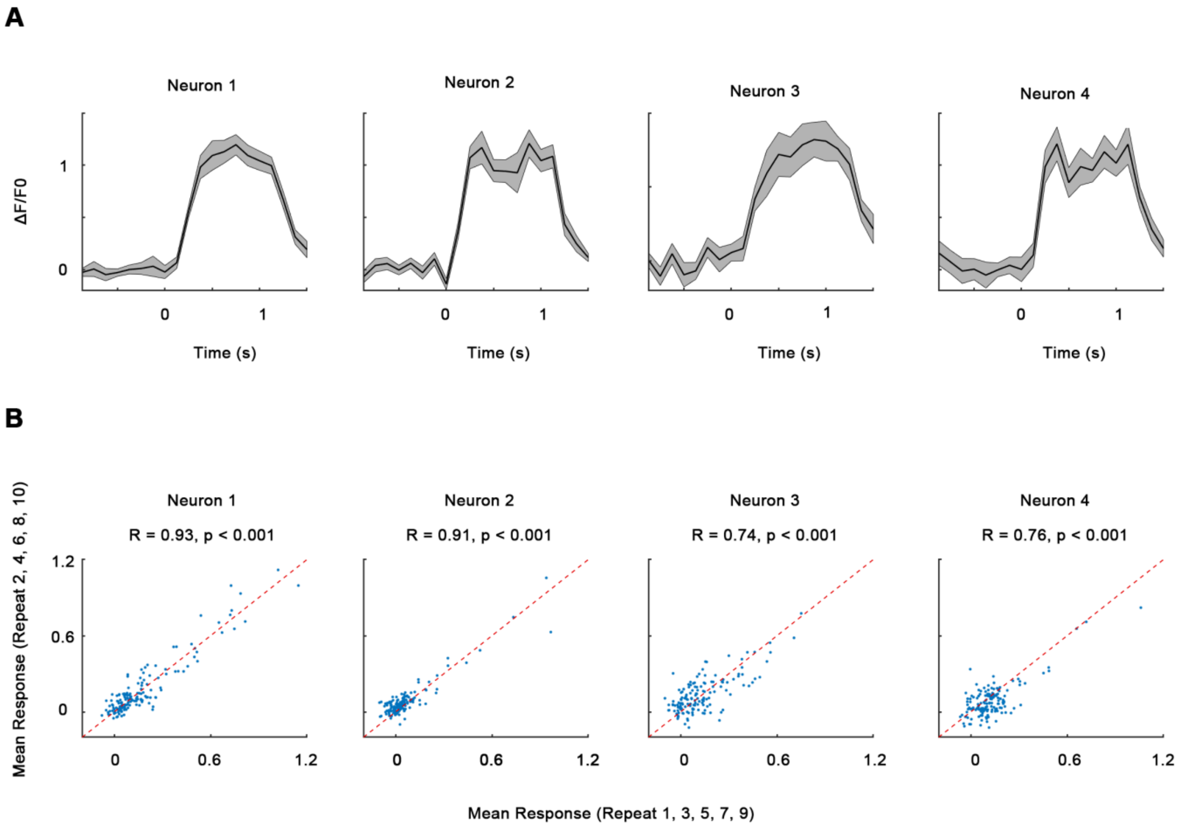


**Figure S2. Related to Figure 2.** Single neuron responses. (**A**) GCaMP timecourses (ΔF/F0, Mean ± SE) of the neurons in Figure 2A-D, each under its optimal stimulus. Trials were synchronized so that 0 sec indicates the time of stimulus onset. (**B**) Scatterplots showing average neuronal responses to the stimuli (bars, curves and corners) in the odd repeats (1, 3, 5, 7) and even repeats (2, 4, 6, 8). The red dash line indicates the y = x line. Pearson correlation R > 0.5 for 474 out of total 535 neurons (88.6%).

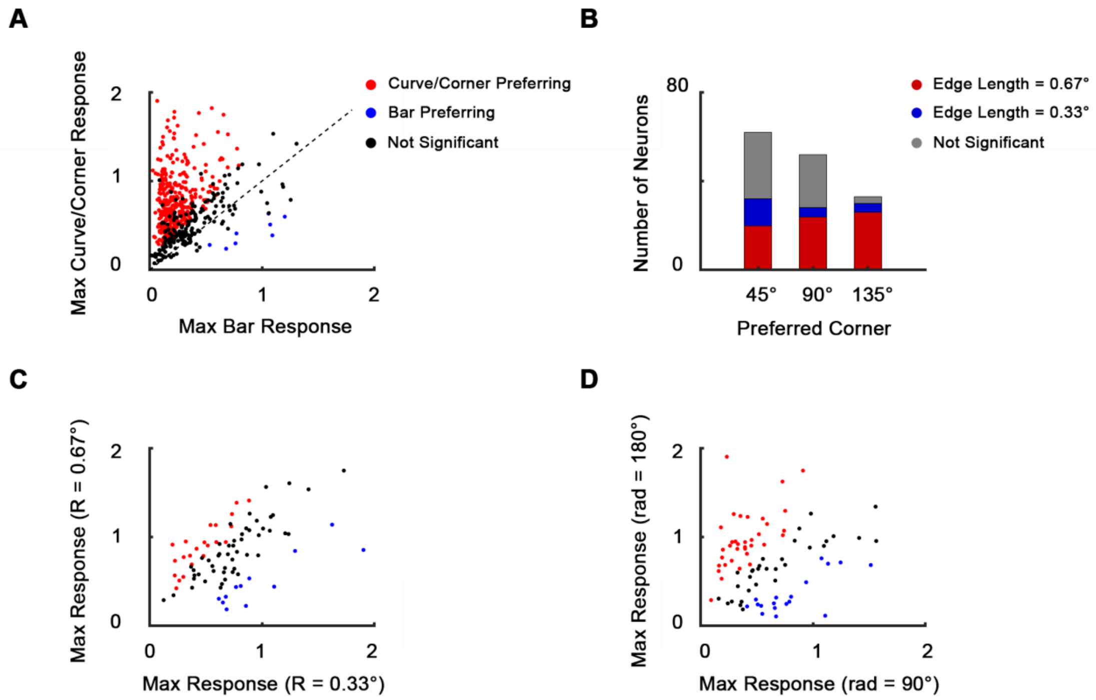


**Figure S3. Related to Figure 3.** Neurons' tuning to curves and corners. (**A**) Scatterplot of neurons' maximum responses to curves and corners against bars. 65.7% of 535 neurons recoded in the imaging area showed significantly stronger responses to curves and corners over bars (n=10, one-way ANOVA, $p < 0.05$). Only 1.5% showed significantly stronger responses to bars over curves and corners. (**B**) Scatterplot of maximal responses to large (R = 20 pixels, 0.67°) against small curves (R = 10 pixels, 0.33°). 84 out of 535 neurons that significantly preferred curves over corners are included, 22 of which significantly preferred big curves and 12 preferred small curves (n=10, one-way ANOVA, $p < 0.05$). 50 neurons were size invariant. (**C**) Scatterplot of neurons' maximum responses to long curves (rad = 180°) against short curves (rad = 120°). Neurons included are the same as B. 35 neurons significantly preferred long curves and 18 preferred small curves. 31 neurons were radian invariant. (**D**) Size and separation angle selectivity of corner selective neurons (147 out of 535 neurons that significantly preferred corners over curves). 62, 52, 33 neurons preferred 45°, 90° and 135° corners respectively. 70 neurons significantly preferred big corners (bar length = 20 pixels, 0.67°), and 20 significantly preferred small corners (bar length = 10 pixels, 0.33°). 57 neurons were size invariant.

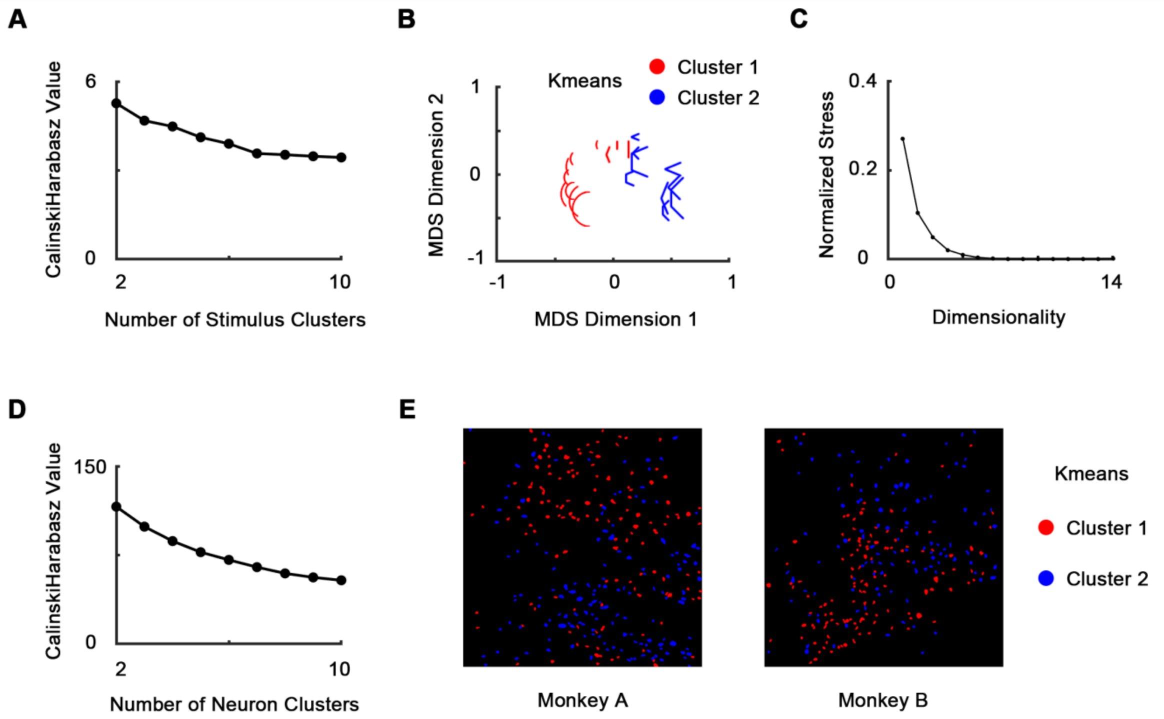


**Figure S4. Related to Figure 4.** K-means clustering analysis. 535 neurons' responses to 20 stimulus forms (2 bars, 8 curves, 6 corners, and 4 Π-shapes, each at 8 orientations) were used. (**A**) We clustered the stimulus forms using population response vector (see methods). Cluster number = 2 according to Calinski-Harabasz criterion. (**B**) The 2 clusters were visualized using MDS. (**C**) The normalized stress of MDS. Stress = 0.104 when dimensionality = 2. (D) We clustered the neurons using neuron response vector (see methods). Cluster number = 2 according to Calinski-Harabasz criterion. (**E**) The two neural clusters are also spatially clustered, and are consistent with CVCNI maps (Figure 3A).

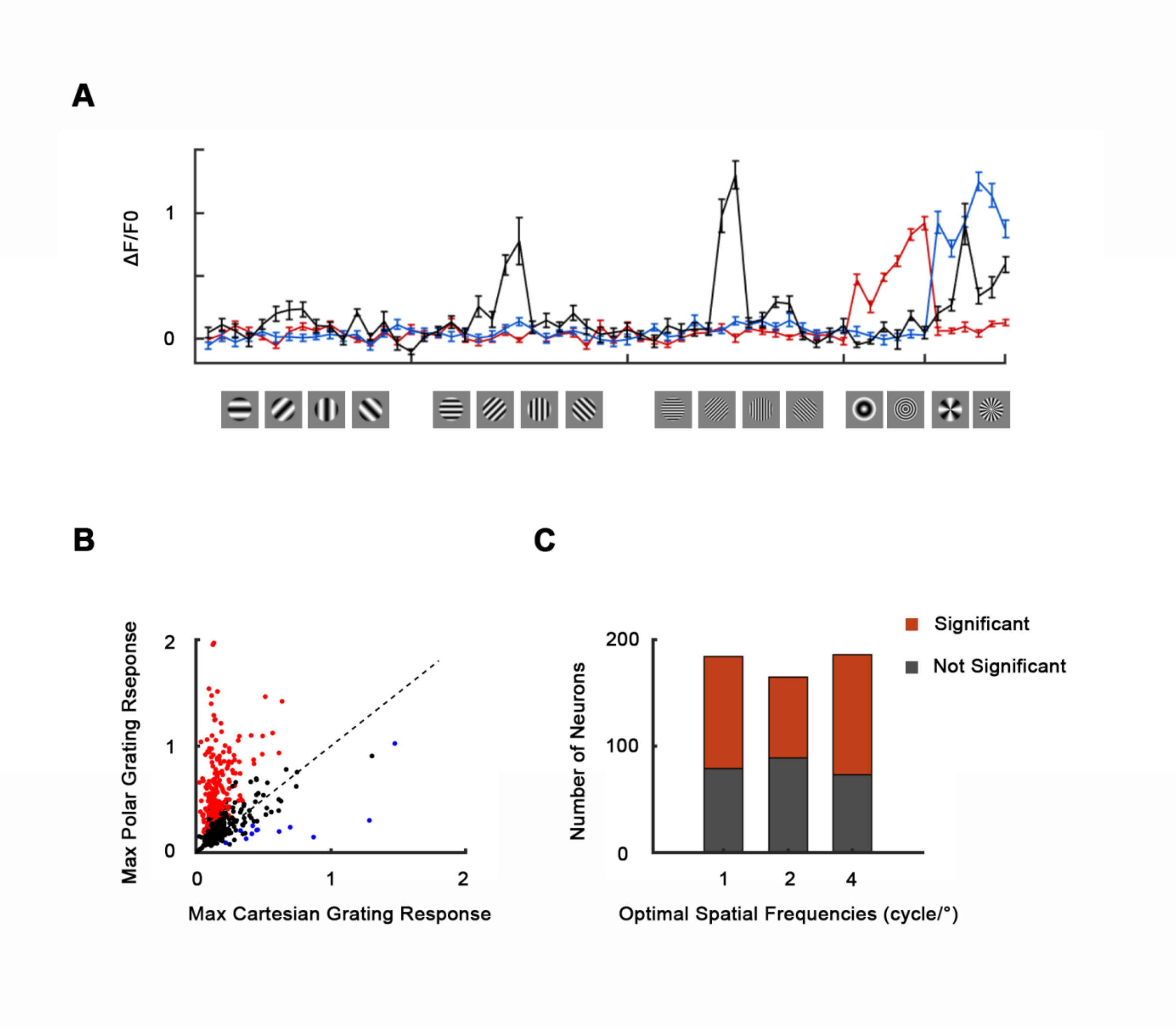


**Figure S5. Related to Figure 5.** Responses to Cartesian, concentric and radial gratings. (**A**) Responses (ΔF/F0, Mean ± SE) of three example neurons to grating stimuli. (**B**) Scatterplot of neurons' maximum responses to concentric and radial gratings against Cartesian gratings. 48.4% of 535 neurons recoded in the imaging area showed significantly stronger responses to concentric or radial gratings (red, $n=10$, one-way ANOVA, $p < 0.05$). Only 2.2% showed significantly stronger responses Cartesian gratings (blue). Black indicates no significant preference ($p >= 0.05$). (**C**) Histogram of neurons' optimal spatial frequencies. Neuronal responses to its optimal gratings (maximum among 10 gratings: concentric, radial and 8 orientated Cartesian gratings) at spatial frequencies of 1, 2 and 4 cycle/° were compared using one-way ANOVA with Bonferroni correction.
